# Distribution and Shared Pathogenicity of Small-Spored *Alternaria* on Solanaceous Crops in Europe

**DOI:** 10.64898/2025.12.11.693629

**Authors:** Gonne Clasen, Zarko Ivanovic, Marta Janiszewska, Remco Stam

## Abstract

Small-spored *Alternaria* species such as *A. alternata* and *A. arborescens* are frequently isolated from diseased potato and tomato plants. However, their respective host ranges and pathogenic behaviours remain poorly resolved, and it is unclear whether their occurrence across hosts reflects true specialisation or ecological opportunism. Limited genetic differentiation among these closely related taxa further complicates their classification as primary necrotrophs or secondary colonisers.

In this study, we analysed natural populations of small-spored *Alternaria* from Germany, Poland, and Serbia using molecular phylogenetics, morphological characterisation, and controlled infection assays to clarify species identity, host associations, and pathogenic potential. Both *A. alternata* and *A. arborescens* were detected across all regions and hosts, indicating broad distribution and ecological overlap. The two species were consistently isolated from foliar lesions and were each capable of causing characteristic *Alternaria* Brown-Spot (ABS) symptoms, thereby fulfilling Koch’s postulates.

Phylogenetic analyses based on *Alt A1* and *RPB2* loci resolved two well-supported species clades and revealed extensive haplotype sharing across more than 1300 km, multiple hosts, and diverse climates, suggesting high gene flow and limited population structure.

The consistent co-occurrence and comparable pathogenicity of *A. alternata* and *A. arborescens* underscore their equal ecological relevance and redefine their roles in ABS epidemiology. These findings indicate that perceived differences in host specialisation may be overstated and that both species contribute equally to *Alternaria* Brown-Spot epidemics in solanaceous crops.

## Introduction

*Alternaria* species are ubiquitous filamentous fungi with a generalist necrotrophic lifestyle, capable of colonising a wide range of hosts and environments. They are of particular agricultural importance as causal agents of early blight on solanaceous crops such as tomato (*S. lycopersicum*) and potato (*S. tuberosum*) (Ivanović et al. 2022; Landschoot et al. 2017a; Schmey et al. 2024). Members of the small-spored Section *Alternaria*, including *A. alternata* and *A. arborescens*, have historically been regarded as secondary colonisers or saprophytes of senescent tissue (Simmons 2000). However, increasing evidence demonstrates that these species can act as primary pathogens, initiating infections independently of large-spored *Alternaria* species (Shi et al. 2023; Sun et al. 2022; Tymon et al. 2016b; Zheng et al. 2015). The disease symptoms are sometimes referred to as *Alternaria* Brown-Spot (ABS), analogous to the diseases caused by small-spored *Alternaria* on other crops, however, some authors cautioned that ABS and Early-Blight symptoms are often not easy to distinguish (Schmey et al. 2024; Vandecasteele et al. 2018).

As generalist necrotrophs, *A. alternata* and *A. arborescens* thrive by exploiting weakened or stressed host tissues (Ramezani et al. 2019), but also persist saprophytically in crop debris, soil, and non-crop hosts (Rotem 1994). This complex lifestyle enhances ecological fitness; saprophytic persistence ensures the accumulation of abundant inoculum, while opportunistic necrotrophy enables rapid colonisation under favourable environmental conditions (DeMers 2022; Thomma 2003). The small, robust conidia of Section *Alternaria* facilitate long-distance dispersal by wind or rain splash (Grewling et al. 2022), with spores surviving a journey over 1,000 km from their source via atmospheric currents (Golan et al. 2023; Plaza et al. 2025). Such persistence and dispersal capacity contribute to the cosmopolitan distribution of small-spored *Alternaria* in global cropping systems (Delgado-Baquerizo et al. 2020; Lawrence et al. 2016).

From a taxonomic perspective, *A. arborescens* remains poorly delimited (Andrew et al. 2009). It belongs to Section *Alternaria*, a morphologically and genetically heterogeneous group in which species boundaries are often indistinct (Lawrence et al. 2016; Woudenberg et al. 2013). Molecular analyses have revealed high sequence similarity between *A. alternata* and *A. arborescens* (Woudenberg et al. 2015) and some authors have proposed that these taxa represent sub-lineages or ecotypes within the *A. alternata* species complex rather than fully distinct species (Andrew et al. 2009; Lawrence et al. 2013; Peever et al. 2004). This taxonomic ambiguity complicates efforts to attribute pathogenicity, virulence, and fungicide resistance traits to specific taxa.

Historically, *A. arborescens* (formerly also *A. alternata* f. sp. *lycopersici*) has been associated primarily with stem canker and fruit-spot syndromes, rather than leaf disease (Simmons 2000; Harteveld et al. 2013; Fontaine et al. 2021). Recent studies (Adhikari et al. 2021; Balodi et al. 2024) found it on leaves too. Field surveys increasingly report the coexistence of *A. alternata* and *A. arborescens* within the same fields and even individual plants (Zheng et al. 2015). Both species have been isolated from tomato and potato, suggesting overlapping ecological niches (Adhikari et al. 2021) and the potential for continuous gene flow among agricultural reservoirs (Ding et al. 2021). This sympatry and niche overlap complicate disease diagnosis and management, as the two species may act independently or synergistically to cause foliar blight symptoms characteristic of *Alternaria* Brown-Spot (ABS) or interact with large-spored *Alternaria* spp. in the Early-Blight Disease-Complex (EBDC) (Landschoot et al. 2017b; Vandecasteele et al. 2018). Despite these observations, the relative abundance and contribution of these organisms to foliar disease development remain poorly resolved at the population level (Salotti et al. 2024).

Both species pose an increasing challenge in agriculture because they tolerate warmer and more *variable* environmental conditions than many other foliar pathogens (Landschoot et al. 2017a; Ščevková et al. 2016). Unlike *Phytophthora infestans*, which is generally recognised as the primary threat in both crops, especially in temperate climates, and which thrives in cool and consistently humid environments, small-spored *Alternaria* can sporulate and infect under warmer temperatures (∼25 °C), intermittent leaf wetness, and fluctuating humidity (Adhikari et al. 2021; Thomidis et al. 2023). As Central European summers become hotter and more irregular, with periods of heat, drought, and sudden rainfall. These conditions are expected to favour *Alternaria* over more climate-sensitive pathogens, potentially increasing its prevalence and impact under future climate scenarios (Delgado-Baquerizo et al. 2020; Fagodiya et al. 2022).

Although no sexual stage has been observed, *A. alternata* and *A. arborescens* maintain high genotypic diversity, providing the evolutionary flexibility needed to persist under changing environmental and agricultural conditions. Population-level studies have reported balanced mating-type ratios and linkage equilibrium (Adhikari et al. 2019; Schmey et al. 2025), consistent with cryptic recombination and the possibility of occasional sexual or parasexual exchange (Adhikari et al. 2021; Meng et al. 2015). Such genetic diversity enhances the capacity of populations to overcome host defences and fungicide pressure (McDonald and Linde 2002; Pereira et al. 2020; Yang et al. 2018). Accordingly, resistance to both quinone outside inhibitors (QoIs) (Grasso et al. 2006) and succinate dehydrogenase inhibitors (SDHIs) (Avenot et al. 2019) has evolved independently in *A. alternata* and *A. arborescens*, underscoring their potential for rapid, parallel adaptation (Avenot and Michailides 2010; Wang et al. 2023).

A deeper understanding of population structure and evolutionary dynamics is essential to disentangle the epidemiology and adaptive potential of these fungi. Small-spored *Alternaria* populations are said to exhibit a predominantly clonal structure interspersed with recombination, facilitating both the rapid spread of successful genotypes and the emergence of novel variants (McDonald and Linde 2002; Meng et al. 2015). High gene flow between agricultural and wild hosts, combined with efficient long-distance dispersal, likely contributes to the regional and transboundary spread of adaptive alleles, including mutations conferring fungicide resistance. Clarifying how *A. alternata* and *A. arborescens* are structured across landscapes and hosts is therefore critical for predicting disease dynamics, managing resistance evolution, and designing sustainable crop protection strategies.

The present study characterises the population composition and pathogenic potential of isolates of *A. alternata* and *A. arborescens* collected from potato and tomato fields across three European countries. Specifically, we quantify their relative frequency and distribution, assess their ability to cause *Alternaria* Brown-Spot, and explore their coexistence as pathogens in solanaceous cropping systems.

## Materials and Methods

### Collection

We collected symptomatic plant tissue from three sites in Germany (July 2023), 14 sites in Serbia (September 2023), and one site in Poland (July 2024) (Figure 1). At each site, two sets of 10 leaves were sampled, with sets separated by at least 50 m to reduce the probability of collecting clonal isolates. We collected one leaf per plant, and leaves were selected based on the presence of typical symptoms consistent with *Alternaria* infection, which could range from small brown spots (i.e., characteristic ABS symptoms) to larger concentric lesions (i.e., those more often associated with EBCD).

**Figure 1.**
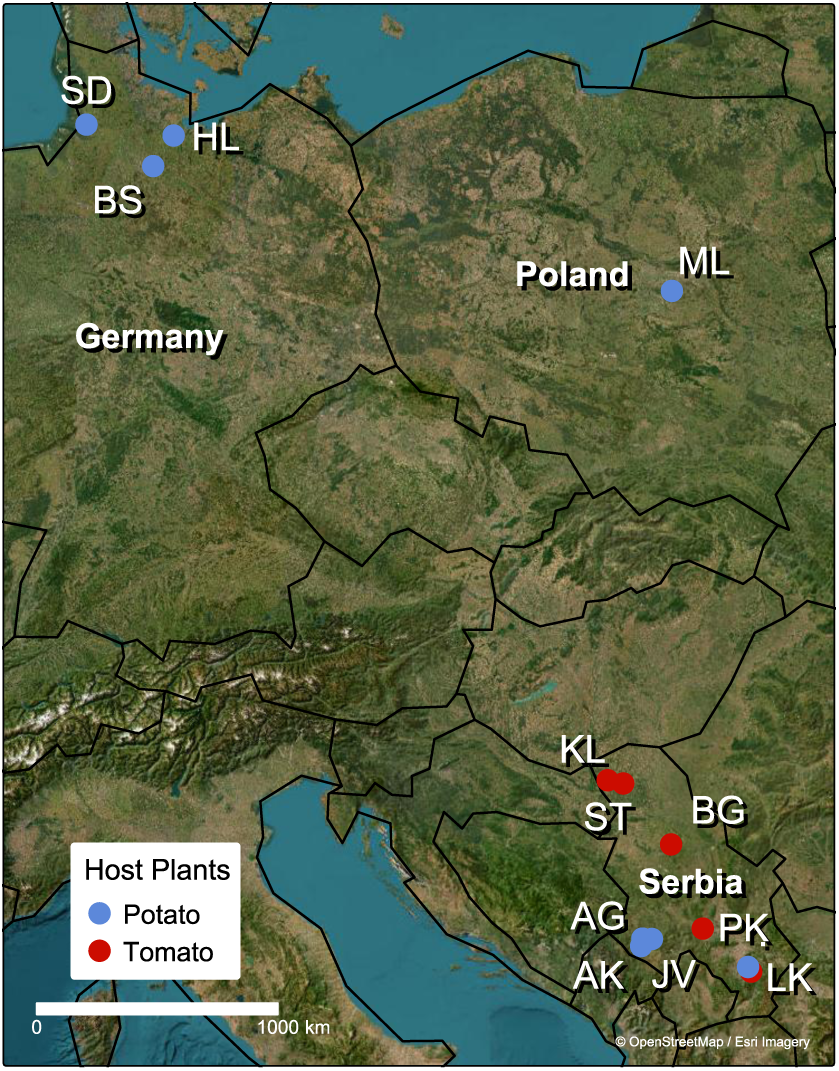
Map of sampling regions in Germany, Poland, and Serbia. Dots are colour-coded by host plant and named after the nearest town.

In Germany and Poland, samples were obtained exclusively from potato (*S. tuberosum*) fields, while in Serbia, we sampled seven potato and seven tomato (*S. lycopersicum*) fields across eight locations. Each isolate was labelled with a standardised code reflecting host species, country, sampling site, and field position (Table 1).

**Table 1.**
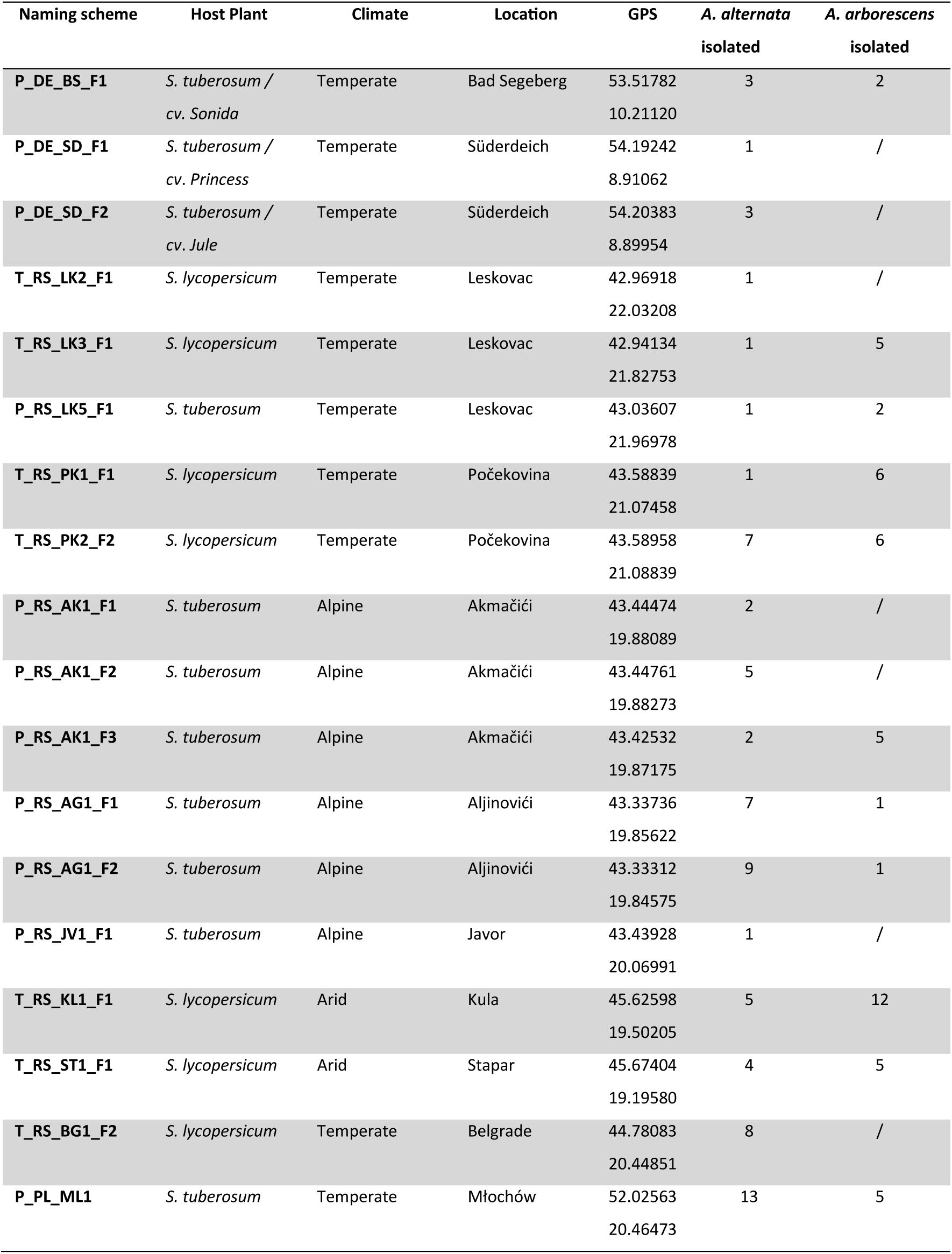
Naming scheme, host plant, climate, location, and GPS coordinates and number of Section Alternaria isolates from sampling sites in Germany, Poland, and Serbia.

### Fungal Isolation, Culture, and Cryopreservation

We established pure fungal isolates by using lesion surface sterilisation and extracting clean inoculum. Dried lesions were submerged in 2 % sodium hypochlorite for 5 min, rinsed in ddH₂O for 3 min, and transferred to Synthetic Nutrient-Poor Agar (SNA) plates (composition: 7.34 mM KH₂PO₄, 9.89 mM KNO₃, 2.03 mM MgSO₄·7H₂O, 6.71 mM KCl, 1.11 mM glucose, 0.58 mM sucrose, 65.41 mM agar, 0.6 mM NaOH) containing 100 ppm Penicillin and Streptomycin to promote fungal growth and suppress bacterial contaminants. Plates were incubated in the dark at 25 °C for 5 days. Under these conditions, we observed growth of small-spored *Alternaria* majorly.

Spores were transferred to Potato Dextrose Agar (PDA) plates (1 g L⁻¹ potato infusion, 5 g L⁻¹ glucose, 3.75 g L⁻¹ agar, pH 5.6 ± 0.2 and partially covered with sterile filter paper to facilitate spore collection. After five additional days of growth, the filter paper was removed, transferred to 2 mL Eppendorf tubes, flash-frozen in liquid nitrogen, and stored at −80 °C for long-term cryopreservation.

### DNA extraction

We generated liquid PDA cultures by autoclaving 200 mL flasks containing 100 mL of tap water supplemented with 97.5 g L⁻¹ PDA. Each flask was inoculated with a 1 mm² piece of spore-covered filter paper and incubated in the dark at 25 °C on a shaker at 120 rpm for 7 days until the onset of melanisation.

Fungal tissue was harvested by centrifugation at 6,000 rpm for 15 min, and 50 mg of fungal tissue was transferred to 2 mL Eppendorf tubes, flash-frozen in liquid nitrogen, and lyophilised (0.01 mbar for 16 h followed by 0.001 mbar for 6 h). Lyophilised material was pulverised using a TissueLyser at 30 Hz for 4 × 30 s intervals.

Genomic DNA was extracted using the Mag-Bind Plant DNA DS Kit (Qiagen, Hilden, Germany) on a KingFisher Flex system (Thermo Fisher Scientific Inc., Waltham, MA) according to the manufacturer’s protocol (Manual Date: May 2024, Revision Number: v7.4).

### PCR and sequencing

Seeing that standard fungal barcode markers such as *ITS* and *TEF1α* fail to reliably distinguish *A. alternata* from closely related species like *A. arborescens* (Woudenberg et al. 2013) we amplified two molecular markers for genes that have been specifically demonstrated to resolve species boundaries within Section *Alternaria* (Woudenberg et al. 2015) and are suitable for phylogenetic analyses: *Alt A1*: primers *Alt-f* / *Alt-r* (Hong et al., 2005a); *RPB2*: primers *RPB2-5F2* / *fRPB2-7cR* (Sung et al., 2007; Liu et al., 2000)

PCRs were performed in 20 µL reactions containing 1 µL Taq polymerase, 5 µL Taq buffer, 5 µL dNTPs (2 mM each), 1 µL forward primer, 1 µL reverse primer, 6 µL ddH₂O, and 1 µL genomic DNA (∼10 ng). Cycling conditions followed the respective published protocols for each marker (Table S1). Amplicons were Sanger-sequenced with the respective forward primers at the Senckenberg Gesellschaft für Naturforschung (Germany).

### Phylogenetic analysis

We processed sequencing data using the AB12PHYLO pipeline (Kaindl et al. 2021) to perform automated quality control, alignment, and phylogenetic inference. Low-quality reads were filtered using Phred ≥30, with 8/10 base trimming at the ends and a maximum low-quality stretch of 5 bases. High-quality sequences were aligned using MAFFT (Katoh et al. 2002) and alignments were trimmed using the Gblocks method in balanced settings (Castresana 2000) to remove ambiguously aligned regions.

Species identification was performed with AB12PHYLO’s integrated BLAST module (Altschul et al. 1990), and results were manually verified against NCBI for top hits. For each locus, Maximum Likelihood (ML) phylogenies were inferred with RAxML-NG (Kozlov et al. 2019) using the GTR substitution model, 1,000 bootstrap replicates, and 20 random + 20 parsimony-based starting trees. Transfer bootstrap expectations (TBE) were used as node support metrics (Lemoine et al. 2018), with blue nodes denoting support values ≥ 70 % and red nodes indicating support values < 70 %. Branch lengths are proportional to the number of nucleotide substitutions per site, representing the amount of genetic change along each branch.

Phylogenetic trees were rooted with *Alternaria solani* as an outgroup, leveraging its larger genetic distance to Section *Alternaria* (Woudenberg et al. 2014). Multigene phylogenetic inference was performed using concatenated loci within AB12PHYLO to confirm species assignments and assess the relationships of *A. alternata* and *A. arborescens* isolates.

### Pathogen distribution and diversity

Each isolate was assigned to categorical *variables* describing *species*, *host plant*, and *climate* based on sample identifiers. To visualise associations among these factors, we generated alluvial diagrams using the ggalluvial package in R (Brunson and Read 2023). Separate plots were produced to highlight species-, host-, and climate-driven flow patterns. Species-host-climate interactions were further summarised using proportional and count-based barplots.

To statistically assess species distribution patterns, contingency tables were constructed for *Species × Host* and *Species × Climate* combinations. Chi-square tests of independence (McHugh 2013) were used to assess whether species frequencies differed significantly between hosts and climate zones. Standardised residuals were computed and visualised as heatmaps to identify over- or underrepresented species-climate combinations. In cases with low expected counts, Fisher’s exact test was considered. Finally, a three-way log-linear model (*Species × Host × Climate*) was fitted using the MASS package (Chambers et al. 2002) to test for higher-order interactions between factors.

Nucleotide diversity (π) and Tajimás D were calculated following (Nei and Li 1979) using the AB12PHYLO alignments.

A Minimum Spanning Network (MSN) (Bandelt et al. 1999) was generated for each species in PopART Version 1.7, using default settings (Leigh and Bryant 2015), based on alignments produced by AB12phylo of the *Alt A1* locus, to visualise haplotype relationships and their association with host plants and sampling regions.

### Morphological characterization

All isolates were grown on SNA plates at 25 °C in the dark for 5 days. Conidia were scraped with a scalpel and suspended in sterile water for slide-mounted imaging. For direct visualisation of conidial chains, spore structures were lifted using transparent adhesive tape and mounted onto microscope slides for imaging. Microscopy was conducted using a ZEISS Primo Star iLED microscope at 400× magnification to capture single conidia. Images of sporulating structures and conidial chains were acquired at 90× magnification using an Olympus SZX12 stereo microscope equipped with a Sony α6400 digital camera.

### Infection assays

Detached leaf assays were performed to evaluate the pathogenic potential of representative *Alternaria* isolates on potato. The potato cultivar used for the assays was *Solanum tuberosum cv. Princess*, originating from the German sampling site in Süderdeich. Tubers were vernalized for four weeks at 5 °C, and plants were grown from tubers and maintained in the greenhouse facility of the Department of Phytopathology and Crop Protection, Institute of Phytopathology, Faculty of Agricultural and Nutritional Sciences, Christian-Albrechts-University, Kiel, Germany. The tomato cultivar used was *Solanum lycopersicum cv. C32*.

Plants were cultivated under standard greenhouse conditions (16 h light / 8 h dark) with supplemental lighting and fertilisation via the irrigation system (1% Sagaphos Blue, Germany). Plants were routinely inspected to ensure they were free of viral infections.

To standardise leaf age and minimise variation, fully expanded leaves from five-week-old potato plants from both cultivars were selected and randomised across treatments. Leaves were placed abaxial side up on PMMA trays (50 × 70 cm) lined with six layers of moist tissue paper to maintain high humidity.

Spore suspensions were prepared by culturing isolates on SNA medium for seven days, washing conidia from the colony surface, and adjusting the concentration to 3 × 10⁴ spores mL⁻¹ using a Thoma counting chamber. Tween 20 (1 mL L⁻¹) was added to the suspension to improve droplet spread. Each leaf received a 10 µL droplet of spore suspension on the abaxial surface; ddH₂O served as a negative control, and *S. chilense* LA4107_3 leaves were included as a susceptible reference.

Trays were covered with transparent lids to maintain humidity and incubated in a growth chamber at 25 °C under a 12 h light/12 h dark photoperiod. Infection success was scored after six days based on the presence of necrotic lesions, and infection frequency was calculated as the proportion of successful infections per treatment. Mean infection frequencies across three independent replicates were compiled into an isolate × host matrix. For heatmap generation, values were centred and scaled per host. Hierarchical clustering was performed using complete linkage, and the resulting dendrogram was used to define the row and column arrangement. The scaled matrix was reshaped to long format using tidyr and plotted with ComplexHeatmap (R version 4.4.1; RStudio 224.12.1+563). A diverging colour scale (steelblue-white-brown) was applied to represent variation in infection frequency. Row-side annotations for species identity, host of origin, and climate region were added by merging metadata with the ordered isolate list. Dendrograms based on the concatenated phylogeny of *Alt A1* and *RPB2* were generated using ggtree and aligned with the heatmap.

To evaluate pathogenicity under near-natural conditions, four *S. tuberosum cv. Princess* plants were inoculated after 8 weeks while still in their pots. Each plant was sprayed with 5 mL of a spore suspension (3 × 10⁴ conidia mL⁻¹) of either of two *A. alternata* or *A. arborescens* isolates. Tween 20 (0.02%) was added to the suspension to ensure homogeneous coverage. Immediately after inoculation, plants were enclosed in a plastic bag for 24 h to maintain high humidity. Disease development was assessed over one week, with photographic documentation taken using a Sony α6400 digital camera before inoculation and at the end of the infection period. The same isolates were also tested in parallel in detached-leaf assays to evaluate comparative virulence.

### Scripts and Data availability

Sanger sequence data have been uploaded to ENA and are available under project number PRJEB105068. All scripts used for the analyses are available under https://github.com/PHYTOPatCAU/AlternariaDistributionPathogenicity.

## Results

### Collection

Section *Alternaria* (small-spored) isolates were present at all 18 sampling sites, irrespective of host plant or climate (Figure 1). Isolates were obtained from potato (*S. tuberosum*) and tomato (*S. lycopersicum*).

We successfully isolated 145 single-spore cultures of small-spored *Alternaria* (Section *Alternaria*), with representation across all farm types, climate regions, and host categories. Sampling in arid northern Serbia yielded fewer isolates than in humid southern areas, likely reflecting lower disease pressure under dry conditions and resulting in fewer sampled sites in the region.

### Morphological observations

Morphological observations of all isolates indicate that all species fall within Section *Alternaria* (Figure 2). Conidia were small, ovoid to ellipsoid, and typically formed in chains, a diagnostic trait of Section *Alternaria* (Woudenberg et al. 2015). *Alternaria alternata* and *A. arborescens* displayed highly similar conidial morphology, and while conidia alone are often insufficient for species-level discrimination, colony growth and sporulation patterns in culture supported the phylogeny-based species assignments.

**Figure 2.**
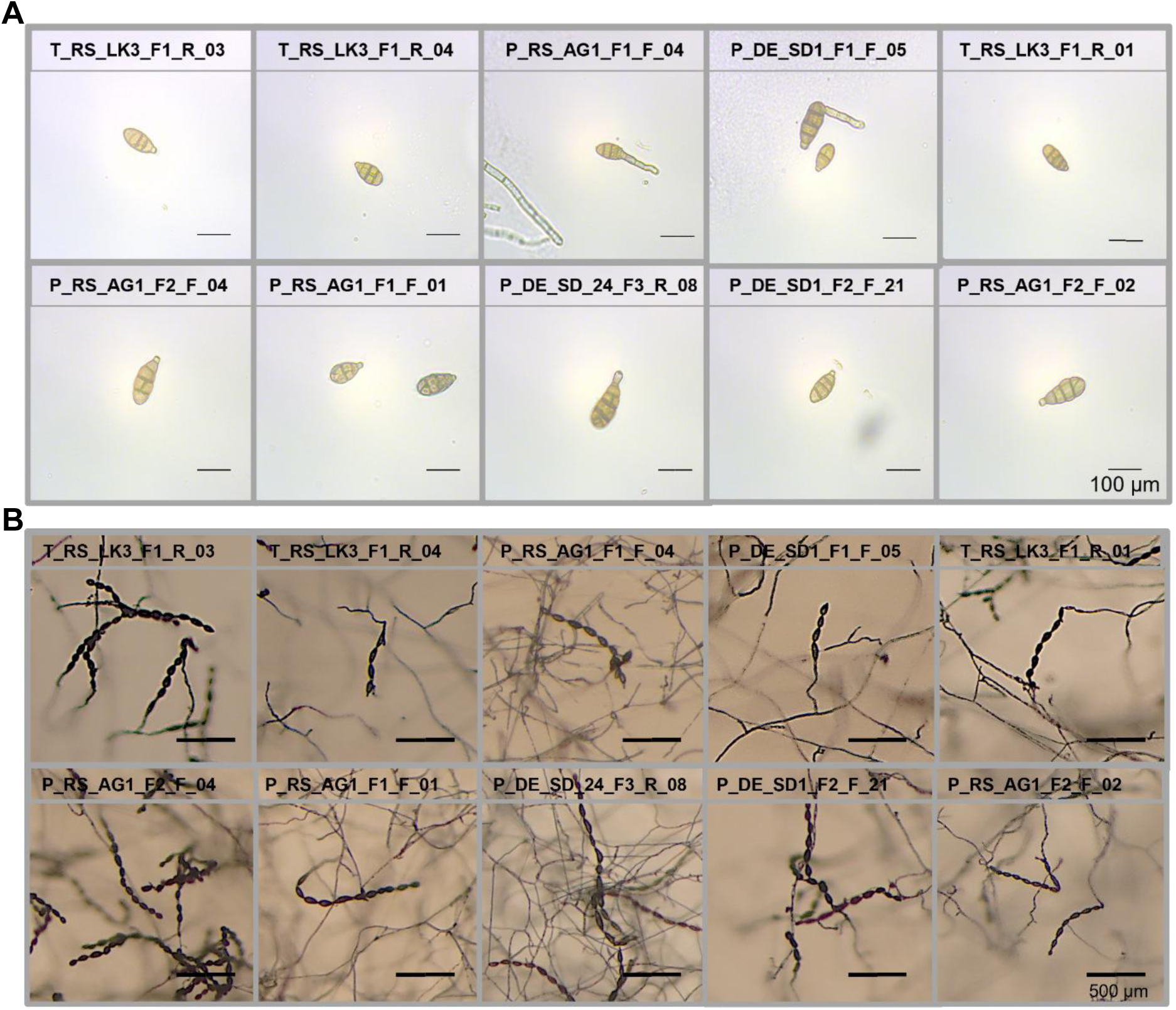
Representative conidial morphology of A. alternata and A. arborescens. The figure shows spores from five A. alternata isolates in the bottom row, and five A. arborescens isolates in the top row, the same isolates used in the infection assays. For each isolate, conidia are displayed both as single spores in water at 400× magnification **(A)** and as conidial chains growing on SNA medium at 90× magnification **(B)**.

Single conidia of *A. alternata* and *A. arborescens* were morphologically indistinguishable. However, their chain-forming sporulation on SNA provided a reliable confirmation of the Section-level identity.

### Molecular identification and phylogeny

We performed molecular species identification using *Alt A1* and *RPB2* as barcode markers and obtained high-quality sequences 145 isolates for both *Alt A1* and *RPB2*. These sequences form the basis for our phylogenetic tree (Fig. 3).

**Figure 3.**
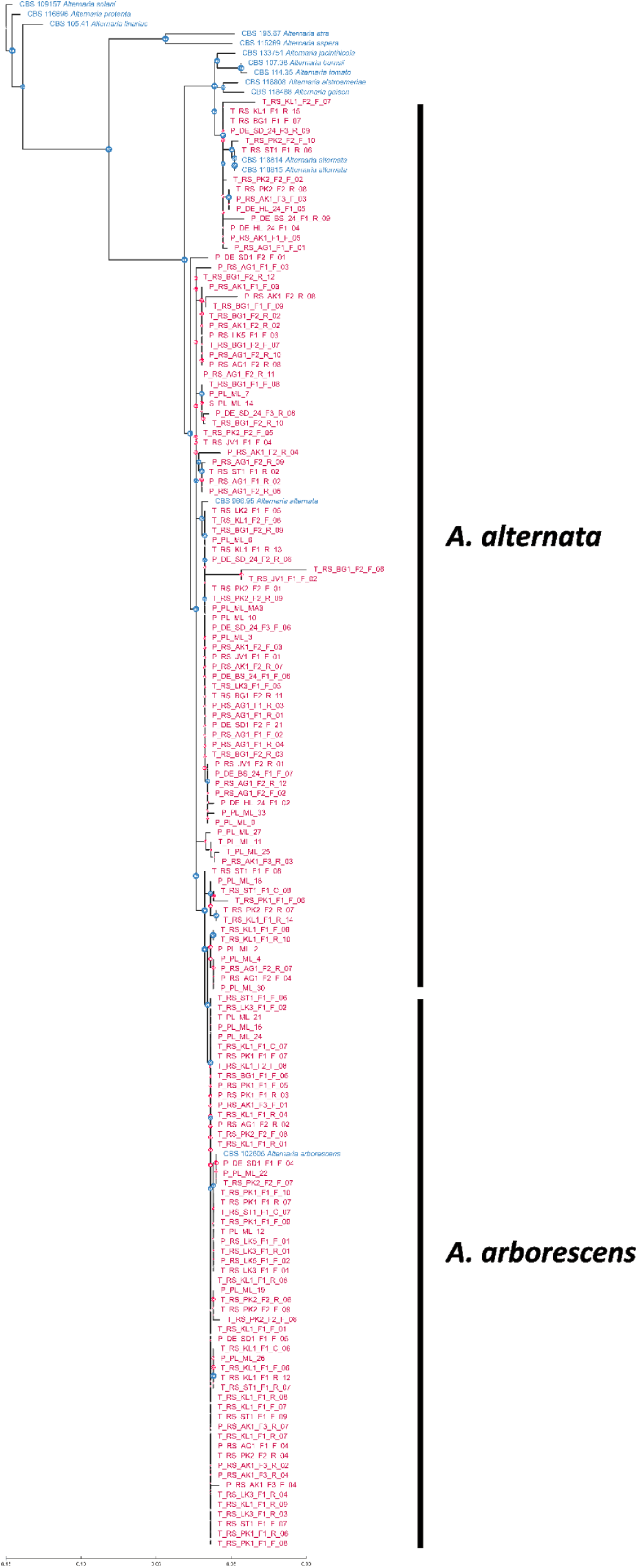
Maximum-likelihood phylogeny generated using AB12PHYLO with the integrated RAxML module from concatenated sequences of two barcode markers, the Alternaria major allergen Alt A1 and the RNA polymerase second-largest subunit RPB2. The tree was rooted to A. solani. Reference isolates included in Woudenberg et al. (2015) are shown in blue, while isolates from the present study are shown in red. Coloured node labels indicate clade support, with blue nodes denoting support values ≥ 70 % (transfer bootstrap expectation; Lemoine et al., 2018) and red nodes indicating support values < 70 %. Branch lengths are proportional to the number of nucleotide substitutions per site, representing the amount of genetic change along each branch.

The concatenated maximum-likelihood phylogeny separated our samples from all large- and small-spored reference species in Woudenberg et al. (2015), except *A. alternata* and *A. arborescens.* Figure 3 shows that all lower terminal branches have very strong bootstrap support (70+) and group with the CBS reference isolate of *Alternaria arborescence* in a monophyletic subclade. Our other samples form polyphyletic groups, almost all of which are closely associated with one of the *A. alternata* reference isolates. We thus conclude that we find both species (and no other small-spored species) present in our sample set.

Within our sample set, *A. alternata* constituted the largest group, with isolates distributed across all sampling regions, farm types, and host plants, whereas *A. arborescens* formed a smaller but clearly defined clade. In all cases, isolates of both species were found within the same fields, highlighting their potential to coexist in the same microhabitat.

We did not find geographic population structure for either species. Isolates from Germany, Poland, and Serbia were interspersed throughout the phylogeny, even within the different *A. alternata* subclades. This pattern is consistent with the high dispersal potential and generalist necrotrophic lifestyle of small-spored *Alternaria* (Schmey et al. 2024).

The single-gene tree for *Alt A1* (Figure S1) was congruent with the concatenated tree’s topology, confirming species placement. Minor discrepancies in the *Alt A1* tree did not affect species recognition or their relative placement in the tree. Together, the single-gene and concatenated phylogenies confirm that all analysed isolates belong to *A. alternata* or *A. arborescens*. These results demonstrate that small-spored *Alternaria* in solanaceous agroecosystems are dominated by these two species, both exhibiting broad ecological range, pathogenicity, and consistent co-occurrence across all surveyed environments.

### Pathogen distribution and diversity

Across all sampling sites, isolates of Section *Alternaria* were consistently recovered, regardless of the host plant or climate region (Fig. 4). Both *A. alternata* and *A. arborescens* were found in Germany, Poland, and Serbia, with both species isolated from potatoes in all countries and from tomatoes in Serbia. This widespread co-occurrence supports the idea that these pathogens are generalists capable of colonising multiple cultivated hosts.

**Figure 4.**
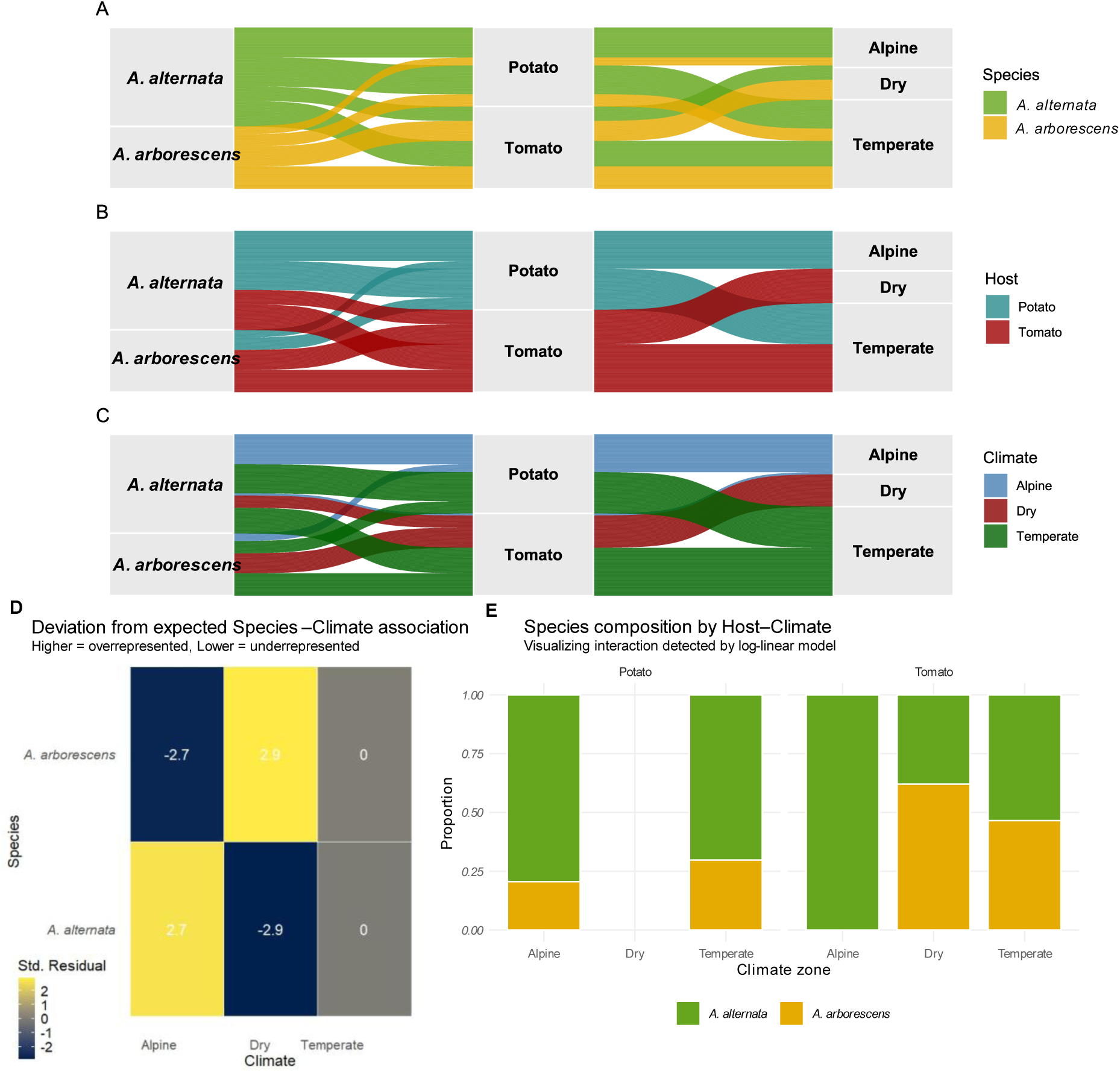
(A-C) Alluvial plots showing the diversity of samples from three perspectives: (A) pathogen species, (B) host plant, and (C) climate region. In each panel, the same alluvial structure is displayed with colours highlighting different variables to emphasise species, host, or climate. **(D)** Heatmap of standardised residuals from the Species × Climate chi-square test, with colour intensity representing deviation from expected counts. **(E)** Proportional species composition derived from a three-way log-linear model (Species × Host × Climate), illustrating the relative abundance of A. alternata and A. arborescens across hosts and climate regions.

**Figure 5.**
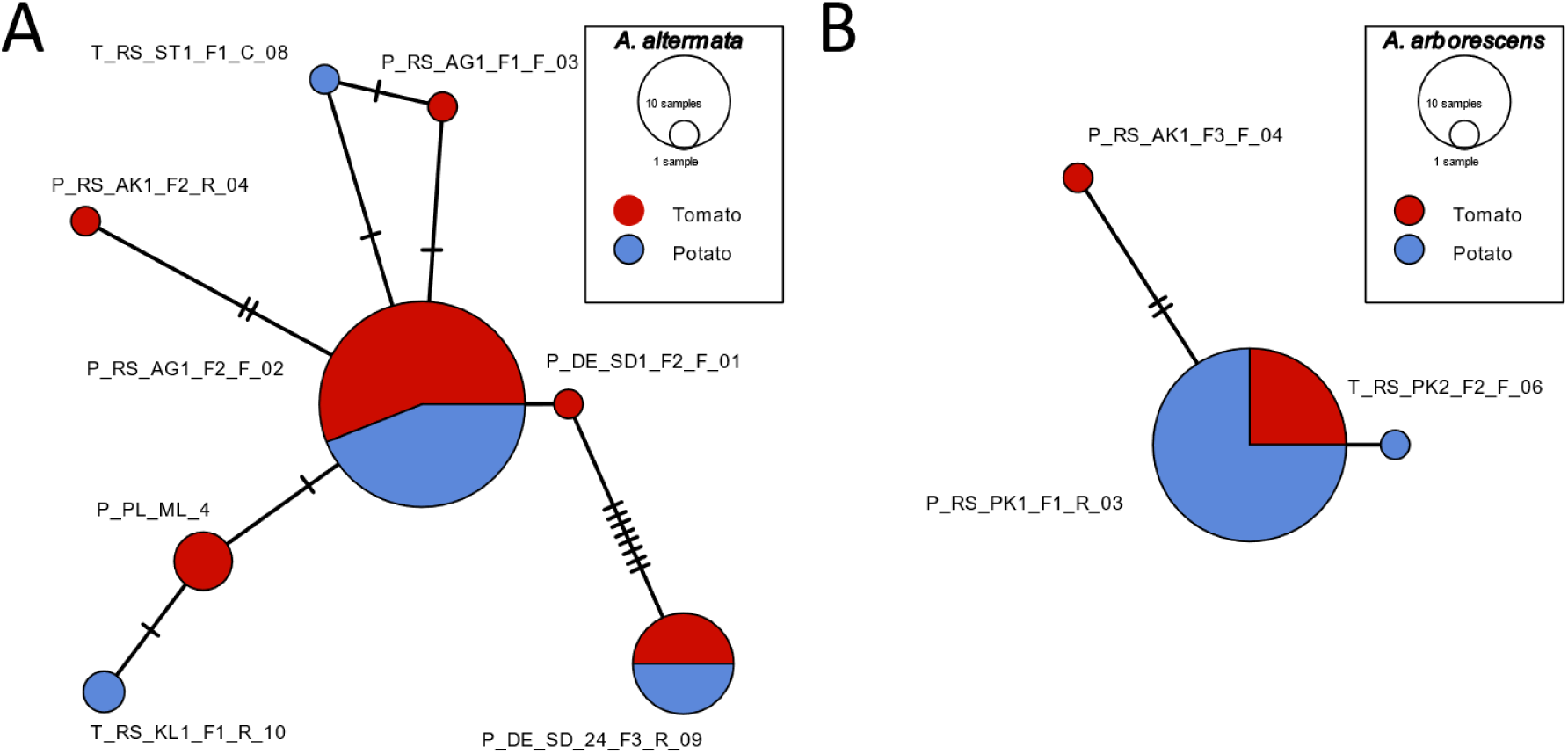
Minimum spanning networks of **(A)** A. alternata and **(B)** A. arborescens based on gene sequences of Alt A1. Each circle represents a haplotype, with circle size proportional to the number of isolates sharing that haplotype. Circles are coloured according to the host plant of origin, and connecting lines indicate mutational steps between haplotypes.

Chi-square tests confirmed that species frequencies differed significantly between hosts (*Species × Host*: χ² = 9.26, df = 1, p = 0.002) and among climate regions (*Species × Climate*: χ² = 12.31, df = 2, p = 0.002). Standardised residuals indicated that *A. alternata* was overrepresented in (colder, wetter) alpine regions (residual = 2.73) but underrepresented in dry regions (residual = −2.90), whereas *A. arborescens* was more frequent in dry zones (residual = 2.90) and less frequent in alpine zones (residual = −2.73). No notable deviations were observed in temperate regions for either species (Fig. 4).

A log-linear analysis of the three-way interaction (*Species × Host × Climate*) further supported these observations. Comparison of an additive model with the saturated interaction model revealed a highly significant improvement in fit when interactions were included (ΔDeviance = 90.98, Δdf = 7, p < 0.001), indicating that host and climate jointly influence species distributions to some extent (Fig. 4).

Overall, these results demonstrate that both small-spored *Alternaria* species co-occur across hosts and climate zones. Yet, subtle climate- and host-associated biases exist, highlighting their ecological flexibility and potential for widespread colonisation in solanaceous cropping systems.

### Genetic diversity and haplotype composition

As expected, our genetic diversity analysis integrated in the AB12PHYLO pipeline, based on the concatenated *Alt A1* and *RPB2* sequences, showed low nucleotide diversity in the markers themselves. Interestingly, multiple haplotypes can be identified within both species. We identified five haplotypes of *A. alternata*, with the dominant haplotype represented by 51 isolates (67 %), and the remaining haplotypes represented by 13, 5, 3, and 4 unique isolates, respectively. Using the same AB12PHYLO pipeline, we identified only three haplotypes of *A. arborescens*, with the dominant haplotype accounting for 39 of 41 isolates.

Our Minimum Spanning Network analysis of the *Alt A1* region across both species (Fig. 5) confirmed that all major haplotype groups occur across countries and host types, reflecting high dispersal potential and a lack of geographic structuring.

**Figure 5.**
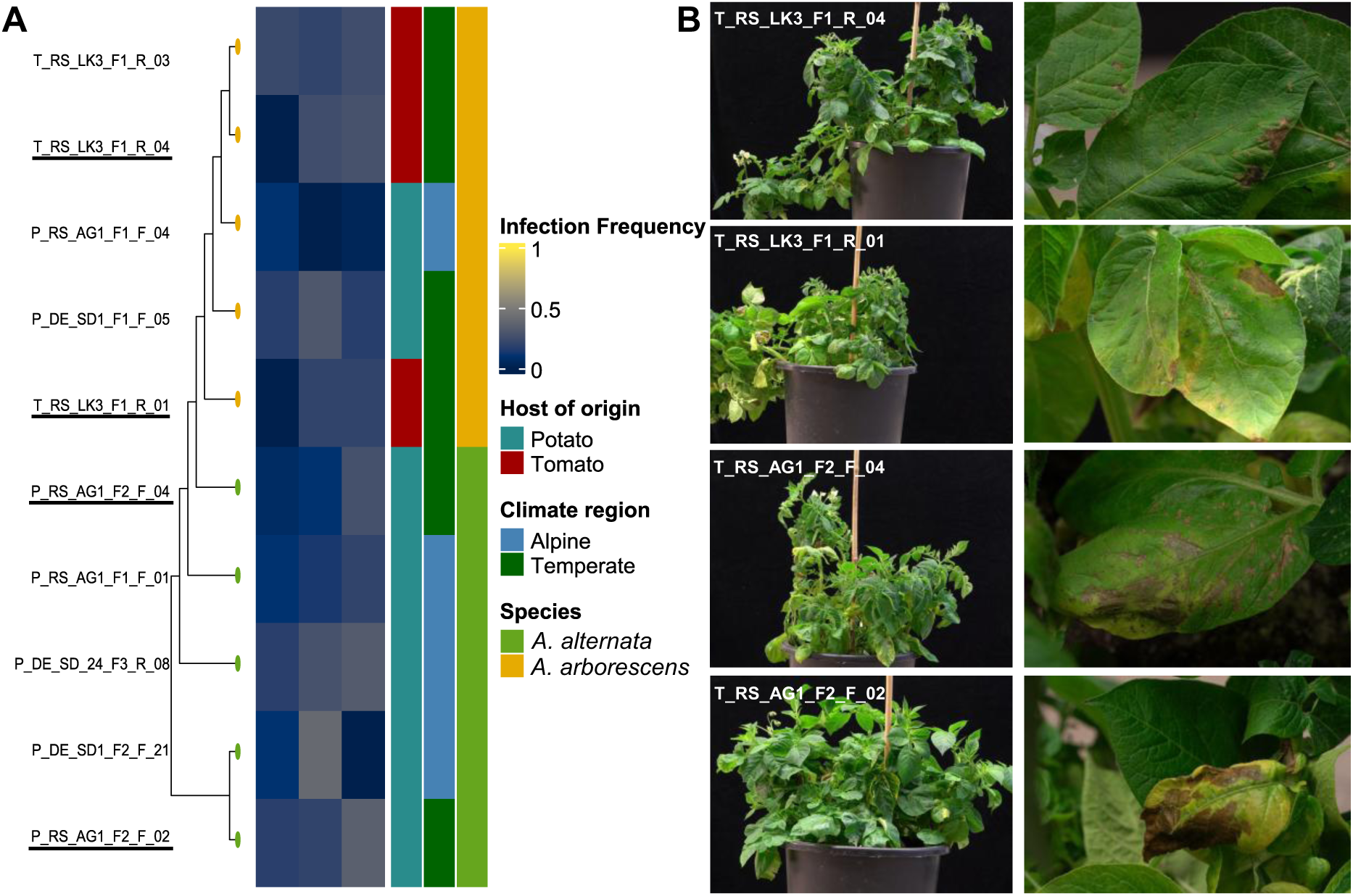
**(A)** Maximum-likelihood phylogeny of all phenotyped isolates paired with a heatmap of infection frequencies on three Solanum hosts. Tip labels correspond to isolate IDs, with tip-point colours indicating species identity. Heatmap cells show infection frequencies (0-1) for each isolate on S. tuberosum cv. Jule, S. tuberosum cv. Princess, and S. chilense LA4107_03. Row annotations denote the host of origin, climate region, and species classification for each isolate. The tree and heatmap are ordered consistently to allow direct comparison of phylogenetic relatedness and pathogenicity profiles. Isolates used for the whole-plant infections are underlined in the tree. **(B)** Whole-plant infection assay validating pathogenicity on S. tuberosum cv. Princess. The left column shows whole-plant images labelled with the respective infecting isolate, and the right column presents corresponding close-up photographs of representative lesions.

Diversity statistics calculated from the concatenated phylogeny showed 71 segregating sites and low nucleotide diversity in both species (Table S2), consistent with clonal reproduction punctuated by occasional recombination. Tajima’s D for both species within Section *Alternaria* was near neutral, which is expected for both *RPB2* and *ALT A1*, with no significant differences between potato- and tomato-derived isolates.

Together, these results indicate that *A. alternata* and *A. arborescens* form a widespread, genetically diverse but weakly structured population, capable of infecting multiple host species within the same agroecosystem. The coexistence of both species across hosts and regions, combined with their broad haplotype overlap, underscores the ecological generalism and dispersal capacity of small-spored *Alternaria*.

### Detached leaf assay and whole-plant infections

Detached leaf assays demonstrated that all of the ten tested isolates were capable of independently infecting potato and tomato leaves, confirming their pathogenic potential on these hosts, and the known *Alternaria*-susceptible tomato species *S. chilense* LA4107_03 (Fig. 5). Across hosts, *A. alternata* infection frequencies varied among infected cultivars, with mean values of 0.15 on *S. lycopersicum cv. C32*, 0.22 on *S. tuberosum cv. Princess’*, and 0.25 on *S. chilense* LA4107_03. Frequencies for *A. arborescens* also varied among infected cultivars, with mean values of 0.06 on *S. lycopersicum cv. C32*, 0.17 on *S. tuberosum cv. Princess,* and 0.23 on *S. chilense LA4107_03*.

The phylogeny-heatmap (Fig. 5A) combines these infection data with isolate metadata and multilocus phylogenetic relationships. The phylogenetic tree, annotated with tip colours signifying species identity, is paired with a heatmap displaying infection frequencies for each phenotyped isolate across the tested *Solanum* hosts. Row-side annotations show the host of origin, climate region, and species classification for each isolate. This visualisation shows that pathogenicity was widely distributed across the phylogeny, with no clade-specific restriction of host-infection ability. Both *A. alternata* and *A. arborescens* isolates exhibited a broad range of infection frequencies, and high- and low-performing isolates were scattered throughout the tree. Row annotations also indicate that the host of origin or climate region did not shape pathogenicity patterns.

To further validate the pathogenic potential observed in detached-leaf assays, we performed whole-plant inoculations on four *S. tuberosum* cv*. Princess* plants, using a subset of four isolates (two *A. alternata* and two *A. arborescens*). After one week, all inoculated plants exhibited characteristic *Alternaria* lesions, confirming that each of the selected isolates was capable of successfully infecting intact plants (Fig. 5B). These results corroborate the detached leaf assay data and demonstrate that both species can cause foliar disease under conditions mimicking natural infection, highlighting their ability to colonise whole hosts rather than only isolated leaf tissue.

These findings provide direct experimental evidence that both *A. alternata* and *A. arborescens* can serve as primary pathogens, challenging their traditional classification as secondary or opportunistic colonisers. The ability of both species to infect multiple *Solanum* hosts under controlled conditions aligns with their more generalist necrotrophic lifestyle and ecological overlap observed in the field.

## Discussion

We consistently recovered small-spored *Alternaria* from symptomatic potato and tomato leaves across all three countries, and both *A. alternata* and *A. arborescens* were repeatedly isolated from the same fields and even from the same host species. This stable co-occurrence across climates, cropping systems, and hosts underlines substantial ecological overlap (Zheng et al. 2015). It suggests that both taxa can contribute to foliar disease pressure in solanaceous crops rather than acting merely as an opportunistic coloniser.

Even though *A. alternata* and species belonging to Section *Alternaria* in general are capable of causing damage to a whole range of other hosts as well, without the need for another primary infection from a species belonging to the large-spored Section *Porri* (Dang et al. 2023; Fan et al. 2023; Shi et al. 2023; Sun et al. 2022), there are still conflicting reports on their virulence in Solanaceae (Tymon et al. 2016a). However, our infection assays confirmed that isolates of both species can infect multiple *Solanum* hosts in the laboratory and cause characteristic *Alternaria* Brown-Spot lesions under glasshouse conditions, demonstrating comparable pathogenic potential under controlled and semi-natural conditions.

The combined epidemiological, phylogenetic, and phenotypic evidence indicates that perceived differences in host specialisation and mode of action between *A. alternata* and *A. arborescens* are possibly overstated. Although *A. alternata* was more frequent overall and showed a modest bias towards cooler, more humid alpine sites, *A. arborescens* was consistently detected at lower frequencies in the same agroecosystems and was enriched in warmer, drier regions, where it nevertheless infected the same hosts. Both species were isolated from symptomatic leaves, and each was capable of initiating disease on healthy plants, contradicting the traditional view of *A. arborescens* as a specific tomato stem pathogen and *A. alternata* as a secondary coloniser, and instead supporting a model of two closely related generalist necrotrophs occupying a shared pathogenic niche.

Genetic data reinforce this interpretation of shared epidemiological roles. Phylogenies based on *Alt A1* and *RPB2* resolved two well-supported species-level clades, with A alternata being polyphyletic as reported before (Schmey et al. 2023; Schmey et al. 2025). Yet, haplotypes of both species were broadly distributed across countries, hosts, and climate zones, with identical haplotypes occurring over more than 1300 km

Looking at the haplotypes only within Section *Alternaria*, we see that our haplotype number is similar to a study from Ding et al. (2019), who used five barcode markers to analyse 40 *A. alternata* isolates. Two of their markers were also used in this study. While Ding et al. (2019) found a mixture of five haplotypes at each of their three sampling locations only about 30 km apart, and only for isolates from potatoes, we found a mixture of our five haplotypes within all sampling regions and host plants (Figure 5) in Germany and Serbia, overall more than 1,300 km apart, and with a 2.5 times higher sample size. Other researchers observed the same trend, with Adhikari et al. (2020), Schmey et al. (2023), and Weir et al. (1998) showing no relationship between the host plant or sampling site and haplotype for *A. alternata.* Blagojević et al. (2020) also found five haplotypes within 27 samples of *A. alternata* found only on rapeseed with low pathogenicity.

Host specificity was not shown to be a factor for the genetic diversity of the large-spored Section *Porri* (Zhao et al. 2023). The host plant or sampling site does not seem to be a factor for isolates belonging to other *Alternaria* Sections either. Kokaeva et al. (2022) failed to show a relationship between phylogenetic groupings and sampling sites. Ozkilinc et al. (2018) report a similar trend. They show a strong clonal structure within Section *Porri* but find no association between a wide range of host plants and the 37 multilocus haplotypes found among isolates from *A. alternata*, as well as the 21 multilocus haplotypes from *A. arborescens.* It is hypothesised that the reproductive mode of *A. alternata* could be the reason for the higher genetic diversity reported in the literature, with Meng et al. (2015) and Stewart et al. (2014) reporting instances of recombination within Section *Alternaria*.

The widespread presence of identical haplotypes across different climate zones and farming systems indicates that both *A. alternata* and *A. arborescens* share a cosmopolitan dispersal ecology. Previous findings of long-distance atmospheric transport of *Alternaria* spores (Golan et al. 2023; Milgroom 2015) would support is. It may facilitate the rapid spread of adaptive genotypes, including those that confer fungicide resistance or increased virulence (Möller et al. 2018). Low nucleotide diversity, near-neutral Tajima’s D, and extensive haplotype sharing indicate a weakly structured, largely panmictic population network, just as shown in Schmey et al. (2025), in which both species disperse efficiently and repeatedly encounter the same host populations, creating ample opportunity for mixed infections and parallel adaptation (Adhikari et al. 2021). However, the use of barcode forms a tremendous caveat in this regard, as they possess general low diversity. Recent whole genome analyses of *A. alternata* show that between isolates genomic presence-absence variation can be observed in genomic regions with possible virulence functions and this variation remained uncaptured in barcode studies.

Pathogenicity profiles mirrored the lack of strong genetic or ecological segregation within our study. High- and low-virulence isolates of both species occurred throughout the phylogeny, and no clade- or host-of-origin-specific restriction of infection ability was observed in detached leaf assays or whole-plant inoculations. Infection frequencies differed quantitatively among hosts and cultivars, and *A. alternata* achieved slightly higher mean infection frequencies than *A. arborescens*. Still, these differences were modest compared with the overall similarity in host range and lesion formation. Together, these patterns argue against strict host specialisation and instead support a scenario where both species behave as flexible generalists that jointly drive ABS epidemics in solanaceous crops.

Given that all haplotypes are generally admixed and that there appear to be no biologically meaningful differences in the ability to infect the tested hosts, one might question whether *A. arborescens* should be recognised as a distinct species or considered part of the larger *A. alternata* species complex, as has been suggested before by others (Fontaine et al. 2021; Harteveld et al. 2013; Simmons 2000).

These findings have several implications for disease diagnosis and management.

First, they suggest that treating Early-Blight symptoms solely as the outcome of infections with large-spored *Alternaria* is likely incorrect. We isolated only a limited number of large-spored isolates from our fields, some of which exhibited apparent EBDC-like symptoms. However, we acknowledge that our methods were designed to isolate only small-spored isolates. Follow-up work would be required to assess the interaction between large and small-spored *Alternaria* in EBDC or ABS complexes. Looking at small-spored species only, we can conclude that treating ABS as the outcome of a single dominant small-spored species is misleading, because mixed populations of *A. alternata* and *A. arborescens* are the rule rather than the exception at the field scale.

Second, the shared ability to infect multiple hosts and the high gene flow across regions imply that adaptive traits such as fungicide resistance or host-specific aggressiveness can arise and spread in either species, potentially moving rapidly through the broader small-spored *Alternaria* complex (Thomma et al. 2003). Consequently, management strategies and monitoring programmes should explicitly account for both taxa, using molecular diagnostics that recognise their close relatedness but avoid assuming distinct ecological roles where the data instead support equal epidemiological relevance.

Finally, this work highlights the limitations of inferring host specialisation and epidemiological dominance from presence-absence data and low-resolution markers alone. While *Alt A1* and *RPB2* clearly separate *A. alternata* from *A. arborescens*, they do not reveal strong host-associated structure or highly divergent pathogenic lineages within either species. Genome-scale data and longitudinal sampling across seasons, hosts, and management regimes will be essential to test whether subtle adaptive differences exist within this shared niche, or whether *A. alternata* and *A. arborescens* are best viewed as two closely related, co-dominant components of a single necrotrophic complex driving *Alternaria* Brown-Spot epidemics in solanaceous cropping systems.

## Contributions

Clasen, Ivanovic, and Stam conceived this study; Clasen, Ivanovic, Janiszewska and Stam planned and performed the experiments. Clasen conducted all analytical steps, produced the figures, and developed the statistical tests. Clasen and Stam wrote the manuscript. All authors read and approved the final manuscript.

## Acknowledgements

We gratefully acknowledge Bettina Bastian, Johanna Preuß, and Maie Bachmann for their invaluable technical support throughout this project. We thank Dr. Severin Einspanier for his insightful feedback and continued technical assistance.

We are grateful to the German farmers from Hof Burmeister and Warfthof Wollatz for granting access to their fields and supporting sample collection. We also extend our thanks to the collaborating farmers and our guides Borko Ivanović from Čačak, Žika Jović from Leskovac and Stanoje Ivanović from Belgrade in Serbia for providing access to sampling sites and local expertise.

We also thank Professor Jadwiga Śliwka for collecting symptomatic leaves in Poland.

Sequencing was performed in collaboration with Prof. Dr. Marco Thines and Dr. Sebastian Ploch.

## Supplementary Tables and Figures

**Table S1.** Primers used for barcode sequencing and PCR conditions.

| Marker | Reference for PCR conditions | Initial denaturation | Denaturation | Annealing | Extension | Final extension | No. of cycles |
| --- | --- | --- | --- | --- | --- | --- | --- |
| ***Alt A1*** | Hong et al. 2005 | 94°C for 1 min | 94°C for 30 s | 57°C for 30 s | 72°C for 1 min | 72°C for 10 min | 30 |
| ***RPB2*** | Sung et al. 2007,<br>Liu et al. 2000 | 95°C for 30 s | 94°C for 15 s | 60°C for 30 s | 68°C for 3 min | 72°C for 7 min | 30 |

**Table S2.** Genetic diversity of the marker genes Alt A1 and RPB2 for both tested species.

|  | *Alt A1* |  | *RPB2* |  |
| --- | --- | --- | --- | --- |
| | $\pi$ | TjD | $\pi$ | TjD |
| ***A. alternata*** | 0.013545 | 1.532110 | 0.008299 | 1.557787 |
| ***A. arborescens*** | 0.000159 | 0.208324 | 0.002023 | 1.620514 |

**Figure S1.**
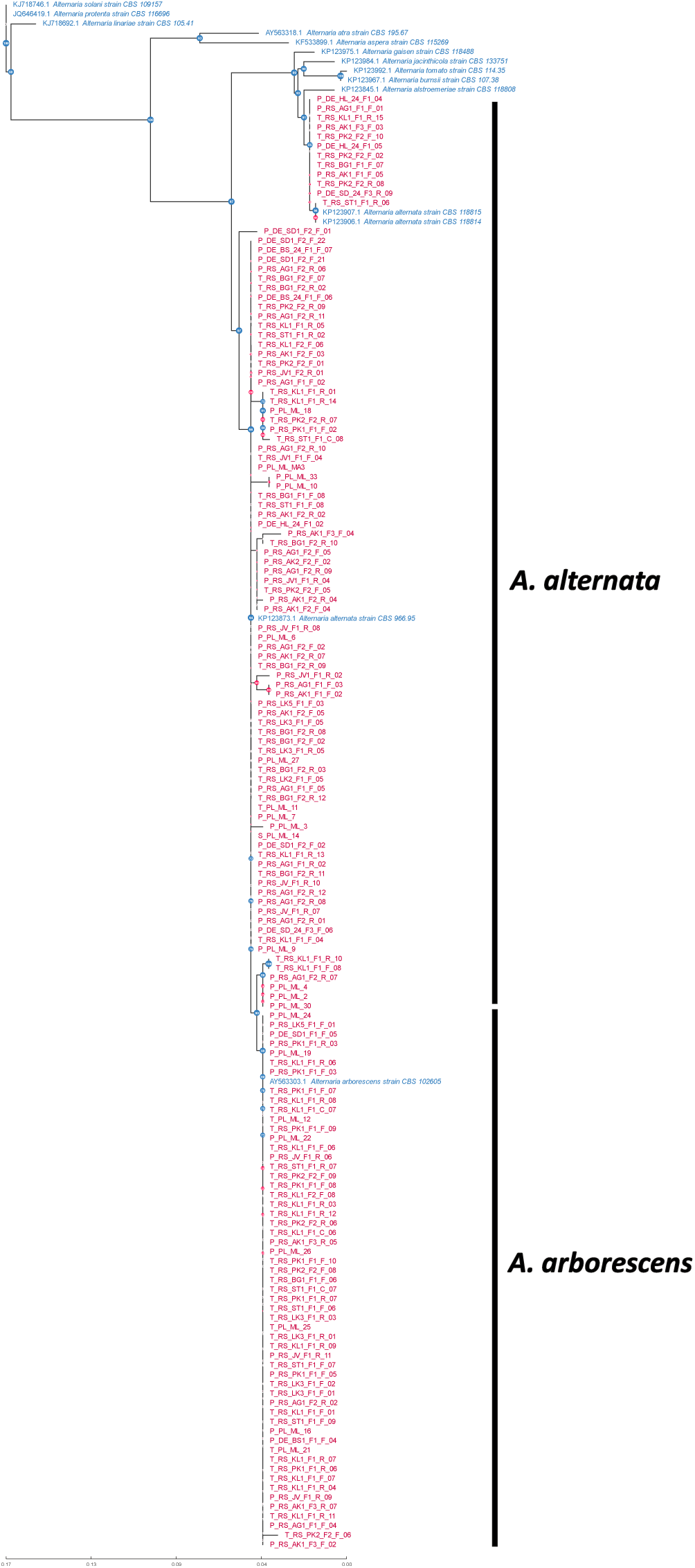
Maximum-likelihood phylogeny generated using AB12PHYLO with the integrated RAxML module from sequences of one barcode marker, the Alternaria major allergen Alt A1. The tree was rooted to A. solani. Reference isolates included in Woudenberg et al. (2015) are shown in blue, while isolates from the present study are shown in red. Coloured node labels indicate clade support, with blue nodes denoting support values ≥ 70 % (transfer bootstrap expectation; Lemoine et al., 2018) and red nodes indicating support values < 70 %. Branch lengths are proportional to the number of nucleotide substitutions per site, representing the amount of genetic change along each branch.

